# Template-Directed RIG-I Agonist Assembly for Targeted Cancer Immunotherapy

**DOI:** 10.1101/2022.12.08.519592

**Authors:** Subrata K. Ghosh, Neil Robertson, Edward Crosier, Michael Dudley, Qiyong P. Liu, Zdravka Medarova

**Affiliations:** TransCode Therapeutics, Inc., 73 Chapel St., Suite B, Newton, MA 02458

## Abstract

Recent developments in the use of pattern recognition receptors (PRRs) aim to harness the innate power of the immune system for cancer therapy. Understanding how to recruit PRRs, such as RIG-I, in a tumor-selective manner is critical for its adoption in the clinic. We describe the use of a tumor-selective template-based agonist of RIG-I to induce type-I IFN signaling and tumor cell apoptosis. The agonist, termed ss-ppp-miRNA-21, comprises a single stranded RNA oligonucleotide modified with a 5’-triphosphate and complementary to an endogenous miRNA enriched in tumor cells. We demonstrate the efficacy of the template-directed approach and detail mechanistic studies validating the hypothesis of a template-directed RIG-I agonist assembly using miRNA-21 as a target. The template-directed strategy described here moves us closer to making RIG-I a clinically relevant target in oncology because it achieves targeted activation of innate immunity in the tumor microenvironment in the context of systemic agonist injection.

## Introduction

Tumor-specific immunotherapy holds significant clinical promise owing to its potential to precisely eliminate malignant tumors without adding toxicity to standard treatments, which makes it attractive as part of the combinatorial therapy for cancers. In recent years, checkpoint inhibitors have demonstrated great success for the treatment of diverse tumor types by releasing the brakes on the immune system. However, they are typically ineffective for patients with ‘cold’ tumors with low levels of tumor-infiltrating lymphocytes (Zappasodi et al., 2018).

Pattern recognition receptors (PPRs), a critical element of mammalian innate immunity, recognize specific evolutionarily conserved structures on pathogens, known as pathogen-associated molecular patterns (PAMPs), resulting in immediate elimination or adaptive immune responses. Upon PAMP engagement, PRRs activate intracellular signaling cascades that eventually cause the release of several proinflammatory cytokines, orchestrating the early host response to infection and laying the groundwork for the subsequent activation and modeling of adaptive immunity (Akira et al., 2006).

Insight into the innate immune response of cells against microbes has inspired the development of cancer immunotherapy based on PPRs, especially retinoic acid-inducible gene I (RIG-I)-like receptors (RLRs), due to the structural simplicity of their ligands, which can be readily synthesized. RLRs are key RNA sensors of virus infection, mediating the transcriptional induction of type I interferons and other genes that collectively establish an antiviral host response. By merely applying an exogenous short 5’ triphosphate double-stranded RNA (5’ppp-dsRNA), a viral infection can be mimicked, allowing direction of the immune response towards otherwise altered or potentially harmful targets such as cancerous cells (Li and Wu, 2021).

Activation of RLRs signaling in the tumor has been found in numerous studies to cause preferential tumor cell death, activate innate immune cells in the tumor microenvironment (TME), and enhance recruitment and cross-priming of adaptive immune effectors, especially in poorly immunogenic, non-T-cell inflamed tumors. These observations have highlighted the possibility to target RLRs for anti-cancer therapy (Rehwinkel and Gack, 2020;McGarry et al., 2021).

Indeed, synthetic RLR mimics are now being studied in preclinical and early clinical trials for the treatment of gliomas, multiple myeloma, breast, pancreatic, and ovarian malignancies (Elion and Cook, 2018;Iurescia et al., 2020). Due to the ubiquitous expression of RIG-I in human cells, it is important to limit its activation in the target site to avoid the side effects of widespread systemic immune activation (Poeck et al., 2008;Ellermeier et al., 2013).

Here, we report a strategy for tumor-selective template-based activation of RIG-I in cancer cells directed by the specific overexpression in tumors of oncogenic miRNA targets. Unlike traditional approaches, which are based on pre-assembled 5’-ppp-dsRNA or 5-ppp-RNA hairpins (Lu et al., 2010, Jiang, 2019 #10), we are using 5’ppp single-stranded RNA which is perfectly complementary to miRNAs known to be enriched in tumor cells (ss-ppp-miRNA). Hybridization of the endogenous target miRNA to the exogenously introduced antisense ss-ppp-miRNA in the cytosol of tumor cells causes the formation of a 5’-ppp-dsRNA, which, as a potent ligand for the cytosolic RIG-I, causes the spatiotemporal activation of the RIG-I signaling pathway preferentially in target cells positive for the endogenous template microRNA. We detail proof-of-mechanism studies using oncogenic miR-21 as a tumor-enriched endogenous template. We show that the proposed strategy not only provides the opportunity for tumor-selective activation of innate immunity through RIG-I agonism but also can be more potent than traditional dsRNA designs.

## Methods and Materials

### Oligonucleotides

The miRNA-21 oligonucleotide, 5’-UAGCUUAUCAGACUGAUGUUGA-3’ (miRNA-21 mimic), and its 100% complement sequence, 5’-UCAACAUCAGUCUGAUAAGCUA-3’ with or without 5’ triphosphate modification (ss-ppp-miRNA-21, ss-miRNA-21), were synthesized by Eurogentec North America (Fremont, CA). A 19-mer 5’ppp-dsRNA positive control for RIG-I activation was obtained from InvivoGen (San Diego, CA; Catalog No. tlrl-3prna).

### Cell culture

The RIG-I reporter cell line HEK-Lucia™ RIG-I cells (Catalog Code, hkl-hrigi, InvivoGen) and the control cell line HEK-Lucia™ Null cells (Catalog No. hkl-null, InvivoGen), and Mouse skin melanoma cells (B16-F10) were cultured in Dulbecco’s Modified Eagles Medium (DMEM, Gibco). All culture media contained 10% fetal bovine serum (serum, Gibco), 100 U/mL of penicillin, and 100 mg/mL of streptomycin. Cells were incubated at 37°C in 5% CO_2_, 5% humidity, and passaged at 2×10^4^ cells/mL when near-confluent monolayers were achieved. Cells were free from Mycoplasma contamination. To generate B16-F10 miRNA-21 transfectants, mature miRNA-21 mimic (0, 0.3, 3, 30, 300, and 1000 ng/mL) was transfected into the cells using LyoVec cationic lipid-based transfection agent (Catalog Code, lyec-12, InvivoGen). Briefly, miRNA-21 mimic was mixed with 100 µL of LyoVec™. The mixture was incubated at 15-25°C for 15 min to 1 h to allow the formation of the complex prior to use for transfection of cells. Ten µL of the LyoVec/miRNA-21 mimic complex was added to 200 µL of culture media.

### Isolation of miRNA and quantitative real-time PCR

After treatment, miRNA was purified from cells using miRNeasy Mini Kit (Catalog No. 217004, Qiagen) according to the protocol recommended by the manufacturer. Then, complementary DNA (cDNA) was synthesized using miRCURY LNA RT Kit (Catalog No. 339340, Qiagen). For each miRNA target, cDNA was diluted prior to use at the ratio of 1:10 with nuclease free water.

The expression analysis of miR-21 was executed by using the miRCURY LNA SYBR Green PCR Kit (Catalog No. 339345, Qiagen) and primers from miRCURY LNA miRNA PCR Assays (Catalog No. 339306, Qiagen) for hsa-miR-21-5p (GeneGlobe ID YP00204230). Reactions were run on StepOnePlus Real-Time PCR System (Applied Biosystems) using the following cycling program: 2 min at 95 °C and 2-step cycling (40 cycles) of denaturation (10 s at 95 °C), and combined annealing/extension (60 s at 56 °C).

The calculation of relative expression was performed using the 2^− ΔΔCt^ method. U6 SNRNA (Catalog No. YP02119464, Qiagen) was used as a reference.

### Western blotting

Cells (3×10^5^) were transfected with 1 µg/mL, ds-ppp-RNA positive control (InvivoGen), ss-ppp-miRNA-21 (8 µg/mL), or ss-miRNA-21 (4 µg/mL) with or without miRNA-21 mimic (1000 ng/mL), using LyoVec transfection agent (Catalog Code: lyec-12, InvivoGen). After a 48-h incubation, the media were aspirated, and the cells were washed twice in ice cold 1x PBS and lysed directly in the plate in IP-lysis buffer (Catalog No. 87787, Pierce/Thermo Scientific) containing Halt Protease and phosphatase inhibitor cocktails (100x) (Catalog No. 78440, Thermo Scientific) for 15□min on ice. Lysates were then centrifuged at 14,000 x g at 4°C for 10 min. The supernatant was collected, and protein concentration was determined using the Quick Start Bradford Protein assay kit (Catalog No. 5000202, Bio-Rad). Equal amounts of the proteins (50 µg) were electrophoresed (125 volts, 1 h) on a 4-20% Mini protean TGX stain-free protein gel (Bio-Rad) and transferred onto nitrocellulose membranes (125 volts, 1 h). Membranes were blocked for 1 h at room temperature with Blocker Blotto in TBS (Catalog No. 37530, ThermoScientific) and incubated at 4°C overnight with primary antibodies in blocking buffer (1:1000 dilution; rabbit monoclonals to RIG-I (Catalog No. 3743), phospho-p65 (Catalog No. 3033), p65 (Catalog No. 8242) and β-actin (Catalog No. 5125) from Cell Signaling Technology, where β-actin served as the reference protein. The membranes were incubated further with anti-rabbit IgG (secondary antiserum), horseradish peroxidase-conjugated secondary antibodies (1:2000 dilution; Catalog No. 7074, Cell Signaling Technology) for 2 h at 37°C. Detection of immunoreactive bands was carried out using Pierce ECL plus western blotting substrate (Catalog No. 32132, Thermo Scientific) and the IBRIGHT CL750 imaging system (Thermo Scientific) according to the manufacturer’s instructions.

### RIG-I activation assay

HEK-Lucia™ RIG-I or HEK-Lucia™ Null cells (1.0, 2.0, 2.5 or 5.0 × 10^4^ cells/well) were seeded in 96-well plates at 70-85% confluence. Then ds-ppp-RNA positive control (1 µg/mL, InvivoGen) or ss-ppp-miRNA-21 (2, 4 or 8 µg/ml) were transfected into the cells using LyoVec™ (Catalog Code, lyec-12, InvivoGen), following the manufacturer’
ss instructions. The transfection was performed immediately after cell seeding. Cells were incubated for 48 h at 37ºC and 5% of CO_2_. Next, 20 µL of culture media was transferred to a 96-well clear-bottom black plate and 50 µL of QUANTI-Luc™ assay solution (Catalog Code: repqlc; Invivogen) was added directly to each well. Luminescence was immediately measured using a Spectramax M3 microplate reader (Molecular Devices) set at a 0.1 second of exposure.

### Cell viability

Cell viability was determined using the CellTiter-Glo Luminescent Cell Viability Assay Kit (Catalog No. G7570, Promega) for each sample according to the manufacturer’s instructions. Briefly, B16-F10 cells were plated in a 96-well clear bottom black plate (Corning, Tewksbury, MA) at a density of 20,000 cells/well in culture media (DMEM supplemented with 10% FBS and penicillin/streptomycin). The ds-ppp-RNA positive control (1 µg/mL, (InvivoGen), ss-ppp-miRNA-21 (2, 4 or 8 µg/ml) or ss-miRNA-21 (2, 4 or 8 µg/ml) were transfected into the cells using LyoVec™ (InvivoGen, Catalog Code. lyec-12, InvivoGen) following the manufacturer’ ss instructions. After a 48 h incubation, 100 µL of CellTiter-Glo (Catalog No. G9242, Promega) was added into each well (containing 100 µL of culture media) and incubated for 10 min at room temperature to stabilize the luminescence signal. Luminescence signal was measured using a SpectraMax M3 microplate reader (Molecular Devices).

### Caspase 3/7 activation

Caspase 3/7 activity was determined using the Caspase-Glo 3/7 Assay Kit (Catalog No. G8091, Promega) for each sample according to the manufacturer’s instructions. Briefly, B16-F10 cells (10,000 or 20,000 cells/well) were seeded in 96-well plates (Corning, Tewksbury, MA) and treated with ds-ppp-RNA positive control (1 µg/mL, (InvivoGen), ss-ppp-miRNA-21 (2, 4 or 8 µg/ml) or ss-miRNA-21 (2, 4 or 8 µg/ml) for 48 h. Plates were allowed to equilibrate to room temperature. One hundred microliters of Caspase-Glo® 3/7 Reagent was added to each well. Plates were incubated at room temperature for 1 h. Luminescence was measured using a SpectraMax M3 microplate reader.

### IFN-_γ_-inducible protein 10 (IP-10/CXCL-10) immunoassay

B16-F10 cells were treated with RIG-I agonist ss-ppp-miRNA-21 and respective controls for 48 h. IP-10/CXCL10 release from mouse cells was assayed using the Quantikine Mouse IP-10 ELISA assay (R&D Systems, Catalog No. DY466-05) using cell-free supernatants of stimulated cells according to the manufacturer’s instructions. The absorbance was measured at 405 nm, and the concentration of IP-10 in the samples was determined by comparison to the standards.

### Testing of ss-ppp-miRNA-21 in a co-culture system

To test for evidence of RIG-I agonism in a co-culture system, we prepared PBMCs freshly isolated from mouse blood using Lymphopure™ (Catalog No. 426201, BioLegend). Blood was diluted 2_∶_1 with PBS and mononuclear cells were separated by density gradient centrifugation. PBMCs were seeded at 100,000 cells/well together with 10,000 B16-F10 cells at an effector□to□target (E:T) ratio of 10:1 in a 96-well plate. Cells were treated with either media as a control, ds-ppp-RNA as positive control (1 µg/mL), ss-miRNA-21 (8 µg/mL), or ss-ppp-miRNA-21 (8µg/mL) and incubated at 37°C, 5% CO_2_ for 48 h. Culture supernatants were evaluated as described above, for IFN-β and IP-10/CXCL-10, produced by PBMCs in response to ss-ppp-miRNA-21 or control constructs.

### Statistical Analysis

Data were expressed as mean ± sem. Statistical comparisons were drawn using a two-tailed t-test. Dose response was analyzed using non-linear regression. All statistical tests were performed using GraphPad Prism software. A *P* value of less than 0.05 was considered statistically significant.

## Results

### Design of the Template-Specific RIG-I Agonist, ss-ppp-miRNA-21

In the present proof-of-concept study, we chose miRNA-21 as a target for template-directed assembly of the RIG-I agonist due to its demonstrated abundance in tumor cells (Bautista-Sanchez et al., 2020). However, the concept is modular and would apply to other miRNA targets enriched in tumor cells. ss-ppp-miRNA-21 is 100% complementary to miRNA-21, modified with a 5’-ppp, and devoid of other modifications (**Fig. 1a**). The proposed mechanism of action of ss-ppp-miRNA-21 is described in Figure **1b** and **1c**. Upon entry into the cell, ss-ppp-miRNA-21 hybridizes to the endogenous miRNA-21 and forms a 5’-ppp-dsRNA (Yoo et al., 2014;Lima et al., 2018). The hybridization event likely results in release of the miRNA from the RISC, as shown previously (De et al., 2013). The 5’-ppp-dsRNA is scavenged by RIG-I in the cytosol causing its activation (**Fig. 1c**). Consistent with the known mechanism of immune modulation through RIG-I agonism (Thoresen et al., 2021), RIG-I signaling promotes rapid rise of type I IFNs and direct cancer cell death, releasing IFNs and pro-inflammatory cytokines as well as tumor antigens (TAs), which triggers cell-mediated immunity (**Fig. 1d**). Both innate immune cells and the adaptive immune system are energized in the remodeled TME. Leukocytes such as NK cells and macrophages boost their cytolytic activity in response to an IFN-rich environment. Importantly, an adaptive immune response is triggered in response to an increase in IFNs and TAs, resulting in the maturation and activation of macrophages and DCs, and enhanced antigen presentation to T-lymphocytes in tumor draining lymph nodes. Naïve T cells are then activated, propagated, differentiated, and transported to the TME, where they can perform an effective anti-tumor response by direct cytolytic activity mediated by perforin and granzymes, as well as indirect cytolytic activity mediated by perforin and granzymes, and the secretion of cytokines like IFNγ and TNFα. Following the death of cancer cells, the majority of tumor-specific effector CD8+ T cells die via apoptosis, but some survive to mature into long-lived protective memory CD8+ T cells (**Fig. 1d**).

**Figure 1.**
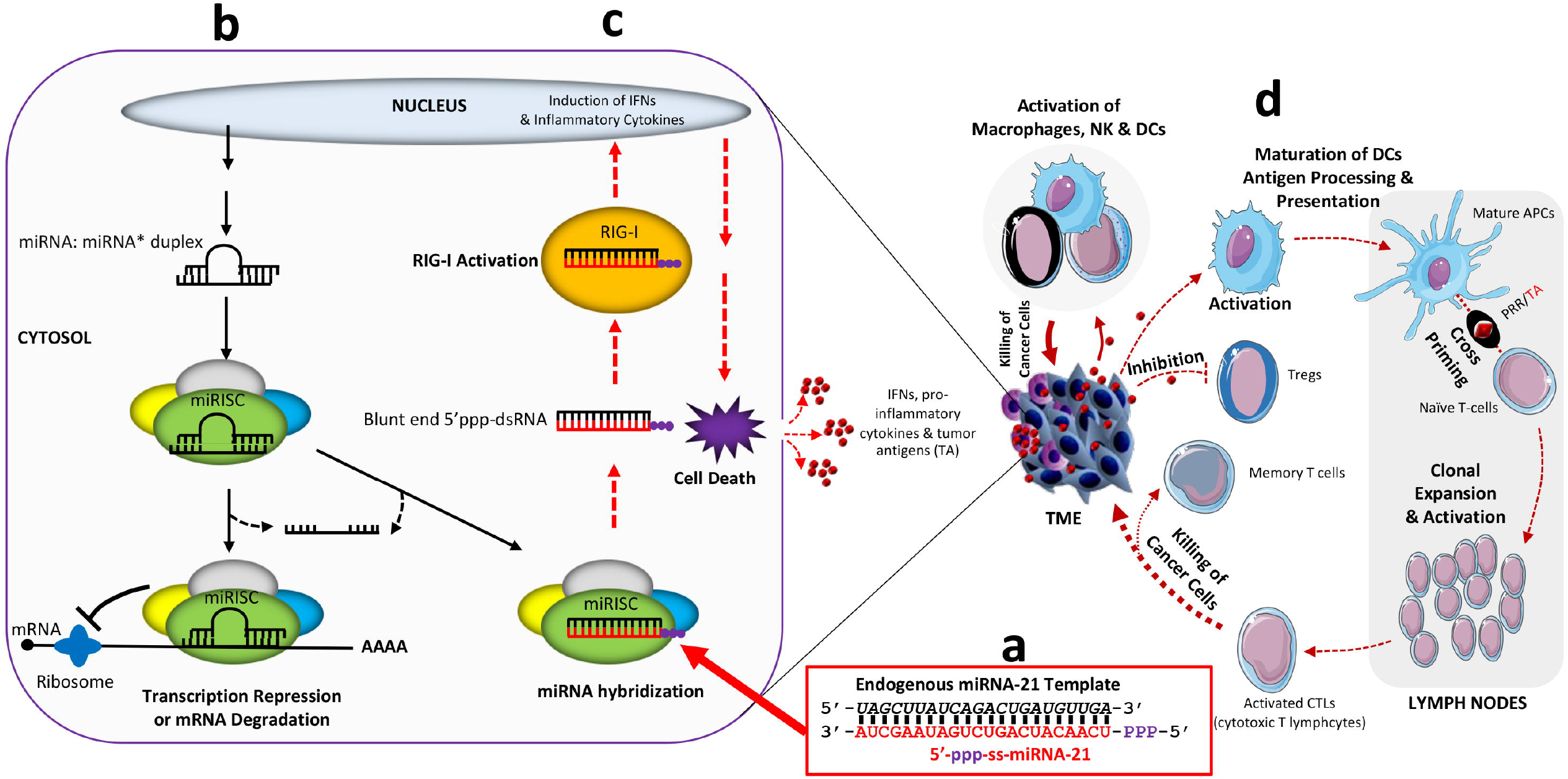
Template-directed RIG-I agonist design and mechanism of action. a) Design of the mir-21-template directed RIG-I agonist, ss-ppp-miRNA-21. The antisense ss-miRNA-21 oligonucleotide, designated as ss-ppp-miRNA-21, was designed to be fully complementary to the target miRNA-21 and to incorporate a 5’ triphosphate modification; b) Biogenesis and function of miRNA (black solid arrow). Starting in the nucleus, imperfectly or perfectly complementary miRNA:miRNA* duplexes are produced in the cytoplasm, where they are incorporated into the Argonaute-containing miRNA induced silencing complex (miRISC), unwound, and the mature miRNA strand retained in miRISC, while the complementary strand is released and degraded. miRISC is guided by the mature miRNA, which directs it to complementary sites in target mRNAs, resulting in translational repression and/or mRNA degradation; c) Activation of RIG-I signaling pathway (red dashed arrow). Upon entry into the cell, ss-ppp-miRNA-21 competes effectively with endogenous mRNA targets (mostly only partially complementary) to bind to the miRNA in the miRISC complex, resulting in release from the RISC and formation of a 5’-ppp blunt ended double stranded RNA. The blunt-ended 5’ppp-dsRNA can be readily captured by RIG-I and trigger its activation. RIG-I signaling in the tumor cells leads to type I IFN-driven immune response and preferential activation of programmed tumor cell death, releasing various immunological factors into the TME; d) Activation of cell-mediated immunity (dark red solid/dashed arrow). RIG-I signaling promotes rapid rise of type I IFNs and direct cancer cell death, releasing IFNs and pro-inflammatory cytokines as well as tumor antigens (TAs). Both innate immune cells and the adaptive immune system are energized in the remodeled TME. Leukocytes such as NK cells and macrophages boost their cytolytic activity in response to an IFN-rich environment. Importantly, an increase in IFNs and TAs triggers an adaptive immune response, resulting in the maturation and activation of macrophages and DCs, and enhanced antigen presentation to T-lymphocytes in tumor draining lymph nodes. Naïve T cells are then activated, propagated, differentiated, and transported to the TME, where they can perform an effective anti-tumor response by direct or indirect cytolytic activity mediated by perforin and granzymes, and the secretion of cytokines like IFNγ and TNFα. Following cancer cell death, the majority of tumor-specific effector CD8+ T cells die via apoptosis, but some survive to mature into long-lived protective memory CD8+ T cells. RIG-I, retinoic acid induced gene 1; IFN, interferon; NK, natural killer; DC, dendritic cell; TME, tumor microenvironment; TA, tumor antigen; APC, antigen-presenting cell; PRR, pattern-recognition receptor; TNFα, tumor necrosis factor alpha; Treg, T-regulatory cells. (Parts of the figure were drawn by using pictures from Servier Medical Art, provided by Servier, licensed under a Creative Commons Attribution 3.0 license).

### The Template-Specific RIG-I Agonist, ss-ppp-miRNA-21, Effectively Agonizes RIG-I and Induces Apoptosis in Melanoma Cells

Our initial feasibility studies focused on the capacity of ss-ppp-miRNA-21 to induce RIG-I activation in the human RIG-I luciferase reporter cell line, HEK-Lucia™ RIG-I. The commercially available cell line stably expresses high levels of human RIG-I and the secreted Lucia luciferase reporter. The reporter gene is under the control of an IFN-inducible ISG54 promoter enhanced by a multimeric IFN-stimulated response elements (ISRE). HEK-Lucia™ RIG-I and HEK-Lucia™ Null control cells can be used to study the role of RIG-I by monitoring IRF-induced Lucia luciferase activity.

We first confirmed high expression of RIG-I in the cells using Western Blot (**Fig. 2a**). We also validated the differential sensitivity of the HEK-Lucia™ RIG-I and HEK-Lucia™ Null control cells to a commercially available traditional RIG-I agonist, consisting of a 5’ triphosphate double-stranded RNA 19-mer (ds-ppp-RNA). There was a highly significant enhancement of luciferase activity in the RIG-I overexpressing cells, as compared to the null cells (**Fig. 2b**).

**Figure 2.**
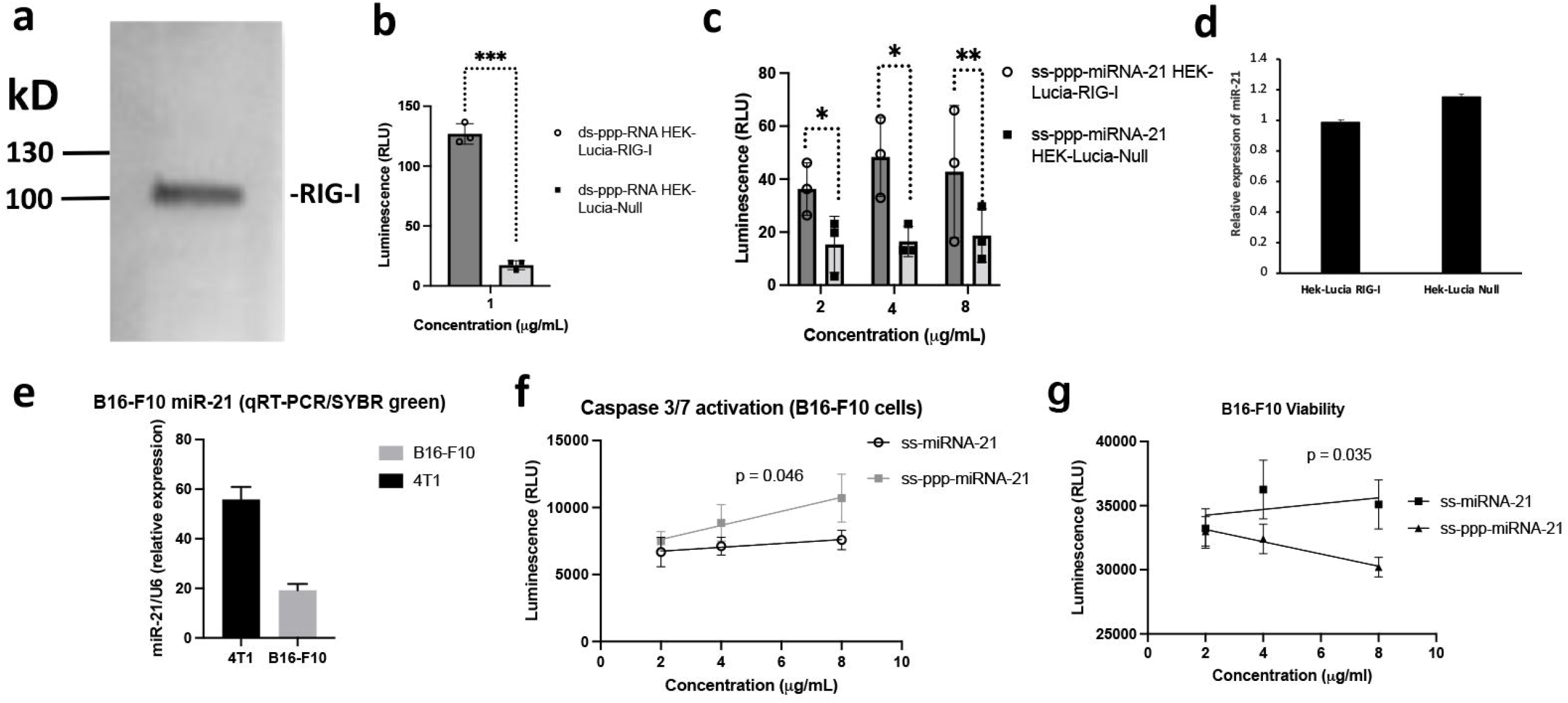
Assessment of RIG-I activation by ppp-ss-miRNA-21. a) RIG-I protein expression in HEK-Lucia™ RIG-I cells. The whole cell lysates were collected and assayed (50 µg) for the RIG-I proteins by Western analysis using specific antibody. HEK-Lucia™ RIG-I cells were found to express high levels of RIG-I protein. b) RIG-I activation by ds-ppp-RNA (+ive control) in HEK-Lucia™ RIG-I and HEK-Lucia™ Null control cells. Cells were transfected with Lyovec control or stated concentrations of ds-ppp-RNA with Lyovec. Luciferase activity was measured in supernatants after 48 h per the manufacturer’s recommendations. The commercially available ds-ppp-RNA was shown to robustly induce RIG-I activation (n = 3; *, p < 0.05; **, p < 0.01). c) ss-ppp-miRNA-21-mediated RIG-I activation in HEK-Lucia™ RIG-I and HEK-Lucia™ Null reporter cell lines. HEK293 RIG-I cells or Null cells were treated with the indicated concentrations of agonists. Luciferase activity was measured in supernatants after 48 h. At all tested concentrations, luciferase activity was significantly higher in the HEK-Lucia™ RIG-I cells than in the HEK-Lucia™ Null cells, indicating RIG-I specific activation (n = 3; *, p < 0.05; **, p < 0.01). d) Relative expression of miRNA-21 in HEK-Lucia™ RIG-I cells and the HEK-Lucia™ Null cells, showing that the two cell lines expressed comparable levels of miR-21. e) Relative expression of miRNA-21 in B16-F10 cells. B16-F10 cells express moderate levels of miR-21, as compared to a reference cell line (murine breast 4t1), known to express abundant miR-21 (n = 3; *, p < 0.05; **, p < 0.01). f) Caspase-3/7 activation (dose dependent) by ppp-ss-miRNA-21 in B16-F10 cells. Cells were transfected with varying concentrations of ppp-ss-miRNA-21 or ss-miRNA-21 with Lyovec. After 48 h cell death was measured by a CaspaseGlo assay. ss-ppp-miRNA-21 induced a dose-dependent increase in caspase activation that was not observed when using ss-miRNA-21 (n = 3, p µ 0.05). g) Cell viability reduction (dose dependent) by ss-ppp-miR21 in B16-F10 cells. Cells were transfected with varying concentrations of ss-ppp-miRNA-21 or ss-miRNA-21. Cell viability was determined using CellTiter-Glo® 2.0 Assay after 48 h of incubation. There was a dose-dependent decrease in tumor cell viability in the presence of ss-ppp-miRNA-21 but not ss-miRNA (n = 3; p µ 0.05).

We next evaluated the capacity of our template-specific RIG-I agonist, ss-ppp-miRNA-21, to activate RIG-I. HEK-Lucia™ RIG-I and HEK-Lucia™ Null control cells were treated with ss-ppp-miRNA-21. We observed significant RIG-I activation in the HEK-Lucia™ RIG-I but not the HEK-Lucia™ Null cells at all three dose levels of ss-ppp-miRNA-21 tested (**Fig. 2c**), despite the equivalent expression of miR-21 in both cell lines (**Fig. 2d**). Given the strict requirement for the formation of an RNA duplex for RIG-I activation (Takahasi et al., 2008;Schmidt et al., 2009;Marq et al., 2010), these results provided support for a template-directed mechanism of RIG-I agonism.

Having established that ss-ppp-miRNA-21 can induce RIG-I in RIG-I overexpressing HEK-Lucia reporter cells, we designed experiments to determine if our template-specific RIG-I agonist can mediate activation of pro-apoptotic signaling in the B16-F10 melanoma cell line. B16-F10 melanoma cells express moderate amounts of miR-21 (**Fig 2e**) and have been used in the past to study intrinsic RIG-I signaling with cell death as an endpoint (Bek et al., 2019). We first measured caspase-3/7 activation in B16-F10 melanoma cells treated with ss-ppp-miRNA-21 or the 5’-ppp-deficient ss-miRNA-21. We observed dose-dependent caspase 3/7 activation that was more pronounced in the presence of a 5’-ppp (**Fig. 2f**). We also found a dose-dependent reduction in tumor cell viability when using the ss-ppp-miRNA-21 RIG-I agonist that was significantly greater than in the 5’-ppp-deficient ss-miRNA-21 (**Fig. 2g**).

### RIG-I Agonism by ss-ppp-miRNA-21 Demonstrates Template Dependence

To further investigate the template-dependence of the observed RIG-I activation when using our ss-ppp-miRNA-21, we repeated the reporter assay using HEK-Lucia™ RIG-I cells transiently transfected with increasing concentrations of a mature miRNA-21 mimic. Transfection with the mimic was carried out prior to addition of ss-ppp-miRNA-21 to avoid annealing of the mimic with ss-ppp-miRNA-21 prior to entry into the cell. We observed a highly significant induction of RIG-I signaling by our ss-ppp-miRNA-21 agonist in cells transfected with 30 and 300 ng/ml of the mimic (**Fig. 3a**), even in cultures of as few as 10,000 cells. The levels of activation with ss-ppp-miRNA-21 were similar to those observed with the commercially available ds-ppp-RNA positive control (**Fig. 3a**). The 5’-ppp-deficient ss-miRNA-21 failed to cause detectable RIG-I activation (**Fig. 3a**). Furthermore, analysis of the dose-dependence of RIG-I activation as a function of miRNA-21 mimic concentration determined an EC50 of 83.4 ng/mL of miRNA-21 mimic when using ss-ppp-miRNA-21. By contrast, the calculated EC50 when using the 5’-ppp-deficient ss-miRNA-21 was 357.9 ng/mL (**Fig. 3b**).

**Figure 3.**
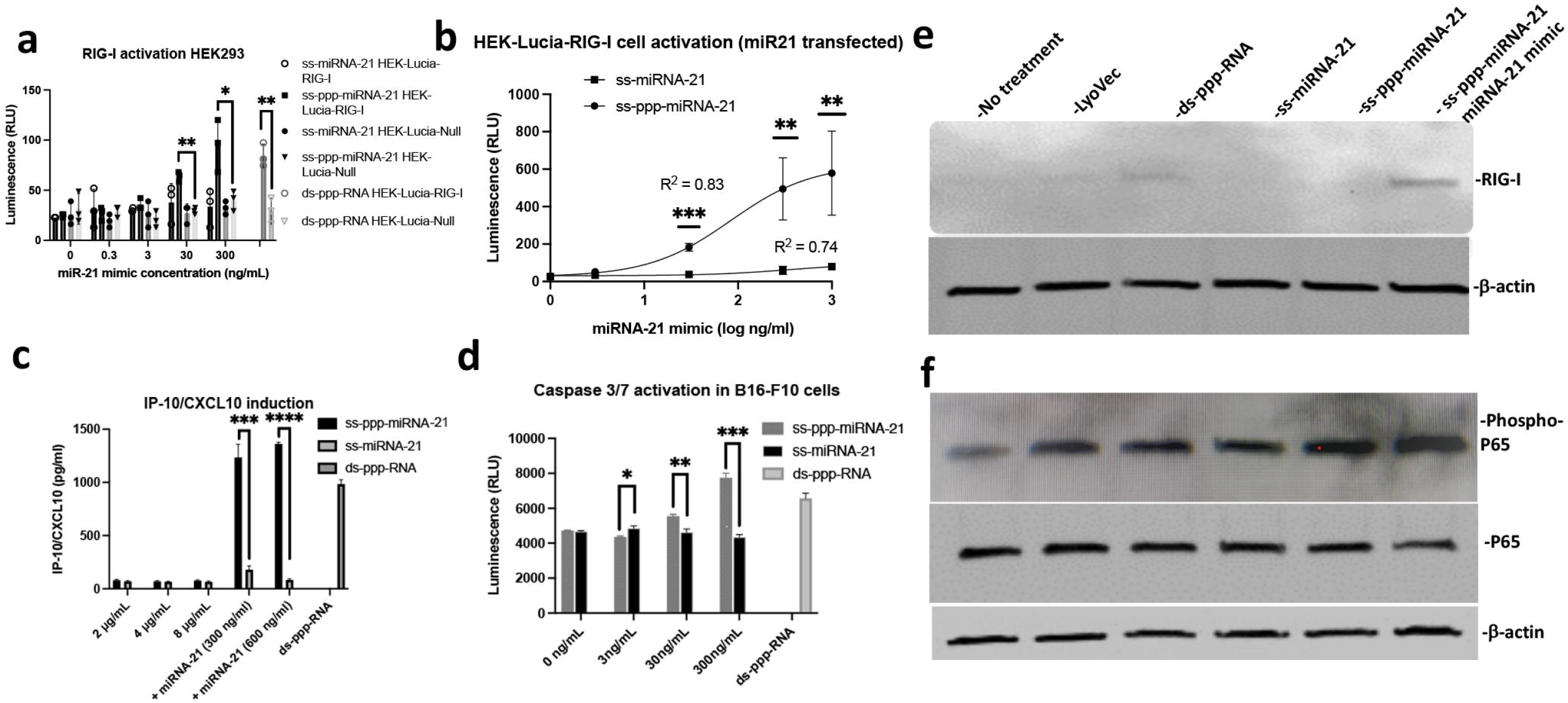
Evaluation of RIG-I activation and related downstream signaling by ss-ppp-miRNA-21 (agonist) and ss-miRNA-21 (control agonist) in the presence of exogenous miRNA-21 mimic. a-b) RIG-I activation by ss-ppp-miRNA-21 in HEK-Lucia™ RIG-I and HEK-Lucia™ Null control cells. Cells were transfected with ss-ppp-miRNA-21 or ss-miRNA-21 (4 µg/mL) along with varying concentrations of miRNA-21 mimic. Luciferase activity was measured in the supernatants after 48 h per the manufacturer’s recommendations. a) There was a dose-dependent RIG-I activation with increasing concentrations of the miR-21 mimic in the in HEK-Lucia™ RIG-I but not the HEK-Lucia™ Null cells, indicating RIG-I-specific activation that is miRNA-21 template dependent (n = 3; *, p < 0.05; **, p < 0.01). b) The response curves to the ss-ppp-miRNA-21 and ss-miRNA-21 resulted in the determination of an EC50 of 83.4 ng/mL for the miRNA-21 mimic when using ss-ppp-miRNA-21. By contrast, the calculated EC50 when using the 5’-ppp-deficient ss-miRNA-21 was 357.9 ng/mL, indicating that the presence of the 5’-ppp rendered the response more potent in this cell line (n = 3; **, p < 0.01; ***, p < 0.001). c) IP-10/CXCL-10 concentration in the supernatants of miRNA-21 transfected B16-F10 cells. B16-F10 cells (with or without transfection with a miRNA-21 mimic) were treated with indicated concentrations of ss-ppp-miRNA-21 or ss-miRNA-21 for 48 h. IP-10/CXCL-10 concentration in the supernatants was quantified by ELISA. There was a significant increase in IP-10/CXCL-10 secretion in cells transfected with the miRNA-21 mimic, when the cells were treated with ss-ppp-miRNA-21 relative to cells treated with ss-miRNA-21 (n = 3; ***, p < 0.001; ****, p < 0.0001). d) Caspase-3/7 activation (dose dependent) in miRNA-21 mimic transfected B16-F10 cells. Cells were treated with ss-ppp-miRNA-21 or ss-miRNA-21 (4ug/mL) along with increasing concentrations of miRNA-21 mimic. After 48 h, cell death was measured by a CaspaseGlo assay. There was a dose-dependent increase in caspase 3/7 activation by ss-ppp-miRNA-21 in cells transfected with increasing concentrations of the miRNA-21 mimic (n = 4; *, p < 0.05; **, p < 0.01; ***, p < 0.001). e and f) western blot analysis of RIG-I; and phosphorylation of NF-kB (p65) protein. B16-F10 cells (with or without transfection with miRNA-21 mimic) were treated with ss-ppp-miRNA-21 (8 µg/mL) and respective controls. Whole cell lysates (50 µg) were analyzed over 8-20% SDS-PAGE and assayed using specific antibodies for indicated proteins. e) There was a moderate increase in RIG-I protein in cells treated with ds-ppp-miRNA-21. Protein induction was most notable in cells transfected with miRNA-21 mimic and treated with ss-ppp-miRNA-21. f) The abundance of phospho-P65 was increased in cells treated with ss-ppp-miRNA-21 relative to control treatments. This effect was seen both in cells transfected with miRNA-21 mimic prior to treatment with ss-ppp-miRNA-21 and cells that had not been transfected with the mimic. No differences in P65 protein were observed, indicating that the effect reflected protein phosphorylation.

Our next set of studies focused on the capacity of the template-specific ss-ppp-miRNA-21 agonist to agonize RIG-I in B16-F10 murine melanoma cells. IP-10/CXCL-10 is a member of the CXC chemokine family and plays an important role in the recruitment of activated T-cells. To confirm IP-10/CXCL-10 expression, IP-10 protein was evaluated in the culture supernatant of B16-F10 cells after treatment with increasing concentration of RIG-I agonist ss-ppp-miRNA-21 and respective controls for 48 hours. As seen in **Figure 3c**, there was robust IP-10 induction in cells transfected with mature miR-21 mimic (300- or 600ng/mL) and treated with ss-ppp-miRNA-21. This effect was not seen with the control treatments or in the absence of the mimic.

Consistent with the known mechanism of apoptosis induction via tumor-cell intrinsic RIG-I signaling in B16-F10 cells transiently transfected with miRNA-21 mimic, we also measured caspase 3/7 activation as a function of miRNA-21 mimic concentration. We found a dose-dependent increase in caspase 3/7 activation which was significantly higher in cells treated with ss-ppp-miRNA-21 as compared to the 5’-ppp-deficient ss-miRNA-21 and comparable to the ds-ppp-RNA positive control (**Fig. 3d**). These data demonstrate that activation of RIG-I using the template-directed approach leads to initiation of caspase-3/7 signaling and induction of apoptosis in B16-F10 cells.

Finally, we analyzed the expression levels of RIG-I, in order to determine if, in addition to RIG-I activation, there was also evidence of RIG-I upregulation in B16-F10 cells treated with ss-ppp-miRNA-21. Low levels of RIG-I were detected in B16-F10 cells. However, in cells transfected with miR-21 and treated with ss-ppp-miRNA-21, there was detectable upregulation of RIG-I (**Fig. 3e**).

One of the mechanisms of immune activation by RIG-I agonism involves activation of the NF-µ B signaling pathway (Ramos and Gale, 2011). In our studies, we analyzed the phosphorylation of the NF-κ B subunit p65 at S536 to measure NF-κ B transactivation. We observed strong phospho-P65 reactivity in lysates from B16-F10 cells treated with ss-ppp-miRNA-21, which was further amplified if the cells were also transfected with a miR-21 mimic. This effect was not associated with increased expression of p65, indicating that it specifically reflected target phosphorylation (**Fig. 3f**). This observation provided further support for effective template-dependent immune stimulation by ss-ppp-miRNA-21.

### ss-ppp-miRNA-21 induces IFN-β and IP-10/CXCL-10 secretion in mouse PBMC/B16-F10 co-cultures

To establish the cell type in mouse PBMCs interacting with the B16-F10 cell line, secretion of IFN-β and IP-10/CXCL-10 was studied in PBMC/B16-F10 co-cultures. The presence of IFN-β in supernatants was detected in ds-ppp-RNA treated co-cultures of PBMC/B16-F10 cells. No detectable levels of IFN-β were observed in co-cultures treated either with ss-miRNA-21 or ss-ppp-miRNA-21 alone (**Fig. 4a**). However, a significant increase in IFN-β secretion was seen in B16-F10/PBMC co-cultures after transfection with mature miR-21 mimic followed by ss-ppp-miRNA-21 treatment.

**Figure 4.**
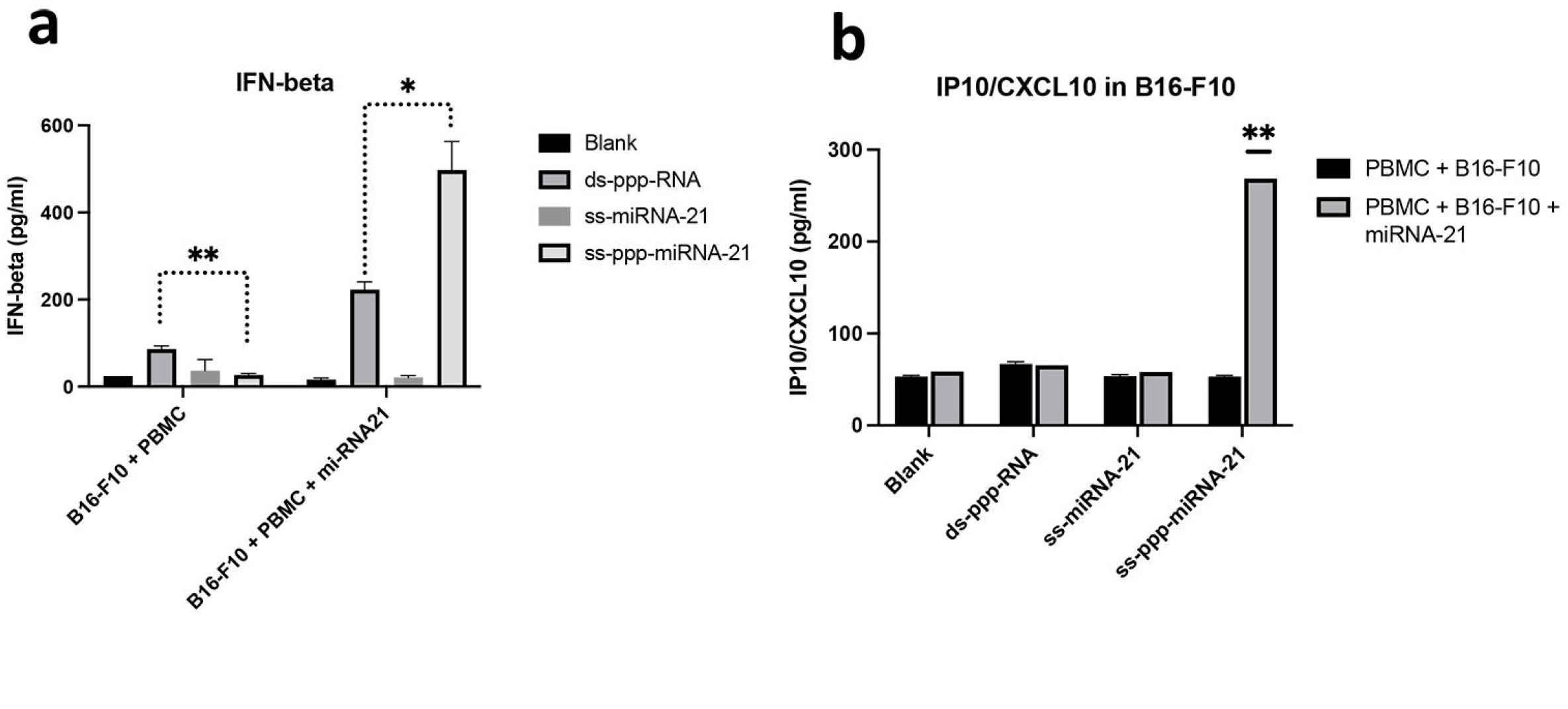
Evaluation of RIG-I activation and related downstream signaling by ss-ppp-miRNA-21 in a co-culture model. a) To evaluate cell cytotoxicity in co-cultures treated with ss-ppp-miRNA-21, we prepared co-cultures of PBMCs freshly isolated from mouse blood and B16-F10 cells at an effectorUtoUtarget (E:T) ratio of 10:1. Cells were treated with either media as a control, or with dsRNA as positive control (1µg/mL), ss-miRNA-21 (8 µg/mL) and ss-ppp-miRNA-21 (8 µg/mL) followed by a 48 h. incubation. Culture supernatants were evaluated for a) IFN-β and b) IP-10/CXCL-10, produced in the co-cultures. Both IFN-β and IP-10/CXCL-10 secretions were significantly induced by ss-ppp-miRNA-21 in the presence of the miRNA-21 mimic relative to controls (n = 2 IFN-β, n = 3, IP-10/CXCL-10; *, p < 0.05; **, p < 0.01).

The supernatants of the above cultures were collected and used to assess the secretion of IP-10/CXCL-10 (**Fig. 4b**). Our results showed that when ds-ppp-RNA or ss-miRNA-21 was tested alone, low IP-10/CXCL-10 secretion was observed. However, in co-cultures of B16-F10 and PBMCs transfected with mature miR-21 mimic and treated with ss-ppp-miRNA-21, there was a dramatic increase in IP-10/CXCL-10 secretion (**Fig. 4b**).

Together these results support the hypothesis that a template-directed approach to RIG-I activation using miRNA-21 as a tumor-enriched target is feasible and potentially associated with more robust responses than the gold-standard dsRNA agonists.

## Discussion

RIG-I detects double-stranded RNA, an RNA virus replication intermediate, and signals through the mitochondrial antiviral signaling protein MAVS, causing type-I interferons to be produced (Akira et al., 2006). RIG-I recognizes viral RNA with an uncapped 5’-di/triphosphate end and a short blunt-ended double-stranded portion, largely independent of the sequence (Brisse and Ly, 2019). These characteristics allow the RIG-I PRR to distinguish self-from non-self RNA and act on divergent viral RNA sequences (Akira et al., 2006).

MicroRNAs (miRNAs) are short noncoding RNAs that fine-tune gene expression at the post-transcriptional level in a spatiotemporal manner. They are typically 20–24 nucleotides long, regulate gene expression by inhibiting protein synthesis or causing mRNA degradation in the 3′UTR of specific genes. The miRNA regulatory network regulates nearly every cellular biological activity, including cell differentiation, death, and proliferation (Lima et al., 2018). The widespread recognition of miRNAs as important biological regulators has increased the demand for strategies to overexpress or inhibit their expression, both to explore function and to treat diseases such as cancer caused by miRNA dysregulation.

Anti-microRNA oligonucleotides (AMOs or anti-miRNA) are antisense steric blocking chemicals that decrease microRNA (miRNA) function by hybridizing and inhibiting the activity of a mature miRNA, resulting in the formation of a miRNA/AMO duplex. Inhibition of the miRNA is accomplished by introducing stabilizing modifications into the AMOs or designing them so that they are composed of LNA/DNA bases (Lima et al., 2018). Alternatively, we have previously shown that when the AMO is a natural ssRNA, the microRNA target is not inhibited, even though there is a transient duplex formation (Yoo et al., 2014). Here, we envisioned that this double-stranded product can be engineered into a potent RIG-I agonist by using a designer AMO with a 5’ppp modification and a 100% sequence complementary to the target oncogenic miRNA.

The success of this approach relies on several key prerequisites. From an oligonucleotide design perspective, it is known that RIG-I binding and activation generally does not tolerate chemical modifications on the ds-RNA agonist (Brisse and Ly, 2019). For that reason, we specifically designed our single-stranded agonist so that it consists of a native RNA oligonucleotide with a phosphodiester backbone in this feasibility study. As shown by us previously, this design does not inhibit the target microRNA but forms a transient duplex with it at a stoichiometry proportional to the abundance of the target miRNA (Yoo et al., 2014). The requirement that the duplex is composed of unmodified RNA bases also poses a limitation on the target template. Templates that are not RNA-based or not linear and capable of forming a blunt-ended duplex with the exogenously introduced ss-ppp-RNA will not be amenable to the design described here (Brisse and Ly, 2019).

Another restriction on the described template-directed design is the requirement of a double-stranded blunt-ended structure for RIG-I activation. It is now known that, even though the presence of a 5’ di/tri-phosphate is not always critical for RIG-I activation (Takahasi et al., 2008;Lu et al., 2010), the availability of a double-stranded structure is essential and, therefore, single-stranded oligos are not capable of RIG-I activation (Takahasi et al., 2008;Schmidt et al., 2009;Marq et al., 2010). Consequently, the potency of ss-ppp-miRNA in terms of RIG-I activation and innate immune activation is critically dependent on its capacity to form a blunt-ended duplex with an endogenous template. For that reason, careful template selection is important to ensure proper hybridization to the exogenous agonist.

Apart from restricting the sequence of the agonist to one that is complementary to an endogenous template, the requirement for a ds-RNA structure for RIG-I activation also provides the unique opportunity to achieve tissue-specific innate immune activation. Current double-stranded or hairpin-based RIG-I agonists are often administered intratumorally (Aznar et al., 2017;Jiang et al., 2019). However, this mode of administration is not clinically feasible, due to limited tissue penetration. Alternatively, systemic administration of a double-stranded RIG-I agonist could trigger systemic activation of innate immunity, due to its lack of tissue selectivity (Poeck et al., 2008;Ellermeier et al., 2013). By contrast, the template-directed approach described here is amenable to safe systemic administration using, for example, nanocarrier delivery vehicles because RIG-I activation will not be triggered in tissues that do not express the endogenous template but will be selective to the target tissue, such as tumors and metastases.

A factor that needs to be considered when designing similar template-directed RIG-I agonists is the abundance of the endogenous template. Based on our studies, efficient RIG-I activation is achieved only when there is a threshold expression of the miRNA template in the target cell. Since miRNA expression is a dynamic process, effective application of the proposed approach for cancer therapy may counterintuitively need to rely on combinations with other therapeutic approaches that upregulate the target oncogenic miRNA. For example, there is evidence that miR-21 is upregulated in response to ionizing radiation (Mahmoudi et al., 2022). It is conceivable that neoadjuvant application of non-ablative ionizing radiation may synergize with the ss-ppp-miRNA-21 RIG-I agonist to produce improved anti-tumor responses.

Finally, for the described approach to be successful, it will be necessary to achieve effective systemic administration through complexation, encapsulation, or conjugation of the ss-ppp-RNA to nanocarriers with optimal PK and tissue distribution profiles. Oligonucleotide delivery to tumor cells using clinically relevant delivery systems remains a challenge but can be overcome as we progress in exploring alternative delivery agents to the traditional lipid nanoparticle or GalNac conjugates (Yigit et al., 2012).

Taking all these factors into consideration, the template-directed approach for RIG-I activation described here holds promise for the development of a novel class of immunotherapeutics for cancer that could rationally complement existing strategies for checkpoint inhibition and neoantigen release to expand the population of cancer patients that can achieve durable disease regressions. The opportunity for systemic, yet tumor selective activation of innate immunity using the template-directed model represents an important step towards effective application of RIG-I and other PRR-agonists for cancer therapy in the clinical setting.

